# Visual shape discrimination in goldfish, modelled with the neural circuitry of optic tectum and torus longitudinalis

**DOI:** 10.1101/2023.06.28.546845

**Authors:** David Northmore

**Affiliations:** Department of Psychological and Brain Sciences University of Delaware

## Abstract

The ability of fish to discriminate shapes visually is well documented. What is not understood are the neural mechanisms employed by fish and other vertebrates that lack a cerebral cortex to distinguish even simple geometric patterns. All the behavioral and anatomical data in fishes point to the midbrain optic tectum as the essential structure, but physiological studies of it have shown only circular contrast-detecting receptive fields and oriented edge detectors with nothing more elaborate that is reliable. Attempting to solve this conundrum, I built a model of tectum with neuron-like elements. When shown objects moving in space, the model forms an attentional locus that tracks one object on a retinotopic layer of simulated tectal pyramidal neurons. The object’s elementary features, when binned together, allow it to be distinguished from other objects to some extent; it fails when objects differ only in the features’ spatial relationships, which fish can use for discrimination. The model’s solution is to bias the attentional locus to the edges of a shape in imitation of goldfish that naturally attend to the top of shapes. Redirection of the attentional locus from a shape’s center is achieved by spatially offsetting synaptic inputs to the pyramidal neurons, effected by the torus longitudinalis (an elongated nucleus at the medial rim of each tectal lobe) and its prolific axonal projections to the pyramidal neurons. The model’s shape discrimination was compared to goldfish in the extensive behavioral data of Bowman & Sutherland (1969) who used shapes with points and projections. One test series showed that fish were sensitive to the relative number of points on the tops of shapes. In another, fish could be trained to discriminate points on the sides. By using different offset connections and only one elementary feature detector for small dark spots, the model successfully emulated the two sets of goldfish data, as judged by significant correlations between model response and fish discrimination over 21 pairs of shapes in each series.

## Introduction

Up until the time of our review of fish vision (Northmore et al., 1978), goldfish and at least 10 other fish species, had been tested for their ability to discriminate geometric shapes by training, mostly in two-choice operant experiments with food reward. The weight of evidence then showed that discrimination behavior was controlled mainly by features such as points, knobs, or corners, especially at the tops of shapes. Also important, but to a lesser degree, were the orientations of principal contours. Other factors such as size, contrast and color were also influential but are of less concern here. The picture that then emerged was of a visual system that recognized a shape by detecting a limited set of features combined with the features’ relative positions on the shape, rather than by constructing a whole-object concept of shapes.

Since then, some fish species have been shown to recognize complex patterns such as the faces of both fish and humans (Parker et al., 2020; Coss & Tyler, 2023; Newport et al., 2016) that would implicate sophisticated discriminative and memory mechanisms. Moreover, goldfish and red-tail splitfins can interpret shape from subjective contours, and in the case of the latter species, recognize a stimulus shape even when partially occluded (Wyzisk & Neumeyer, 2007; Sovrano & Bisazza, 2009). Fish also seem to experience several classic perceptual illusions (Agrillo et al., 2020). Such capabilities in humans and other mammals are typically ascribed to cortical mechanisms; the fact that fish and other “lower” orders without a cerebral cortex exhibit them might suggest the operation of non-telencephalic or proto-cortical brain mechanisms, that with the emergence of mammals were transferred to the cortex (Zhaoping, 2016). Whatever the merit of that idea, the present modelling study proposes a mechanism for shape discrimination by fishes that is centered in the midbrain and is quite different in operation from the hierarchical, feature detecting mechanisms of cortex (Serre, 2014).

How the discrimination of simple geometric shapes might be accounted for by known neural mechanisms in fishes is a mystery. We can largely rule out a significant role of the telencephalon because its ablation in goldfish does not impair the learning of a circle-square discrimination (Flood, 1975) or other kinds of visual discrimination (Bernstein, 1962; Iwai et al., 1970; Ohnishi, 1991). However, ablation of the midbrain optic tectum eliminates all visual abilities (Yager et al., 1977) except for large-field luminance detection (Northmore & Sharma, 1979). The importance of tectum versus telencephalon is further underlined by the fact that the great majority of optic fibers terminate in tectum, forming a detailed retinotopic map in the superficial layers (Meyer, 1980), whereas only small numbers of optic fibers terminate in the several pretectal nuclei (Robles et al., 2014), which are, therefore, unlikely to contribute much to spatial discriminative abilities. While there are pathways that could transmit visual information from tectum to the telencephalon, they are indirect via relatively small thalamic/pretectal nuclei (Ito & Kishida, 1978; Hagio et al., 2018; Yañez et al., 2018). The few functional studies of teleost pretectum have found visually driven units that responded to spot motion with large receptive fields (Rowe & Beauchamp, 1982; Friedlander, 1983; Wang et al. 2020). Visual units in teleost forebrain respond with habituating and temporally complex spike trains (Saidel et al., 2001). None of these studies indicate anything resembling the shape selectivity of neurons in the ventral stream of the mammalian visual cortex (Serre, 2014).

We must, therefore, look to the optic tectum and its neural mechanisms to understand pattern discrimination in fishes. The tectal layers, *stratum opticum* (SO) and *stratum fibrosum et griseum superficiale* (SFGS), receive most of the retinal ganglion cell terminals whose activity can be recorded with extracellular electrodes and distinguished from postsynaptic cell activity by various criteria. The most comprehensive investigations using these methods in goldfish tectum by the Maximovs and their associates revealed distinct functional types found at characteristic depths in SFGS (Maximova et al. 2019). The units encountered most superficially were directionally selective, responding to edges moved in one of three directions. Next in depth were units sensitive to edge orientation, horizontal or vertical. Also at this depth, but less commonly encountered, were units without sustained activity, sensitive to small spots of positive or negative contrast, but insensitive to large field luminance changes. At a deeper level, there were units with larger receptive fields that responded in sustained fashion to dimming or lightening. The latter two classes are important components in the model proposed here.

The visual properties of tectal neurons in teleosts have been difficult to characterize by electrical recordings and have been a disappointment to seekers of feature detectors. Their responses to visual stimuli are variable, often strongly habituating with repetition, and exhibit large, often multi-center receptive fields (Schellart & Spekreijse, 1976; O’Benar, 1976; Guthrie & Banks, 1978; Sajovic & Levinthal, 1982; Guthrie & Sharma 1991) that seem ill-suited for analyzing the details of shapes.

Advanced techniques of recording and imaging activity in larval zebrafish are providing a wealth of new detail on tectal processing. The data so far that is relevant to shape recognition include demonstrations of selectivity for stimulus orientation (Nikolaou et al., 2012) and size. Small and large stimuli differentially activate populations of superficial inhibitory neurons (SINs) with somas at the tectal surface and neurites at various depths in the retinorecipient neuropil (SO & SFGS) that engage with periventricular neurons, possibly sharpening their size-selectivity (Del Bene et al., 2010; Preuss et al., 2014).

Despite these advances, we still have no evidence that tectal neurons elaborate shape analyzers. A 60-year absence of such evidence is beginning to look like evidence of absence. Apparently, the discriminative functions of tectum are achieved in a radically different fashion from mammalian visual cortex. Instead of the multiple retinotopic maps arrayed in separate cortical areas, connected quasi hierarchically (Felleman & Van Essen, 1991), tectum possesses a single retinotopic map composed of several substrata receiving input from different kinds of retinal ganglion cells (Robles et al., 2013). Because most tectal neurons processing this input extend radially through the tectum, it is hard to see how a hierarchical network of any depth could be formed, an architecture considered key to mammalian pattern vision. Understanding that, it is claimed, has been advanced by modelling vision with deep learning in hierarchical convolutional neural networks (Mély & Serre, 2017). While the recognition abilities of these systems are impressive, they differ from brains in their learning algorithms, in their exclusively feedforward architecture and, most notably, in their inability to attend selectively.

### Attention in tectum

It can be argued that the evolution of vision for catching prey immediately necessitated the invention of selective attention. The first jawed fishes (ca. 500 mya) that pursued other mobile animals needed to hold a focus of attention on one of them in order to direct an effective attack. Prey seeking must also have required the development of discriminatory abilities ensuring pursuit of edible species rather than the distasteful or the dangerous. The ability to form an attentional focus for preferentially analyzing a region of the visual field and to allow the focus to track a moving object could then have given rise to a form of object perception in which the constituent features of the object are bound (Triesman & Gelade, 1980; Reichenthal at al., 2020), allowing it to be classified into useful categories such as edible, inedible, noxious, inanimate etc. The requirements for a mechanism to do this can be stated quite simply: (i) a retinotopic representation of the scene by simple analyzers, such as for size, edge orientation and color, (ii) a winner-take-all type of attention-forming focus in retinotopic coordinates, (iii) selection of those features present at the focus, and (iv) classification based on those selected features leading to approach or avoidance behavior.

The idea for a plausible neural implementation of this mechanism comes from a modelling study (Northmore, 2017) that demonstrated how midbrain mechanisms, specifically in the tectum and *torus longitudinalis* (TL) could enable a predator fish to hold a focus of attention on the prey it is chasing despite rapid displacements of the retinal image during saccadic eye movements. The model proposes that the attention-focusing mechanism involves a “pointer network” composed of a numerous cell type in tectum, the pyramidal neuron whose soma lies in SFGS (Type I of Meek & Schellart, 1978) (See Figs. 7 & 8). It receives input from retinal terminals in SO and SFGS and has an extensive dendritic tree in the most superficial tectal layer, the *stratum marginale* (SM). There, it receives excitatory input from the marginal fibers originating in granule cells of the TL, a paired, elongated nucleus attached to the medial edges of the tectal lobes (Vanegas et al., 1979; Wullimann, 1994; Folgueira et al., 2007). The model assumes that pyramidal cells are mutually inhibited via interneurons to make a winner-take-all configuration. One visual stimulus object, the most salient out of many that may be present in the visual field, will then activate a cluster of pyramidal neurons at its retinotopically corresponding position in tectum, forming the attentional focus. Given some additional neural circuit refinements, the focus is able to track the object moving in space. However, an eye saccade shifts the retinal image of the object too rapidly for it to be tracked, and the focus of attention will likely settle on a different target object with unfortunate consequences for hunting. Nevertheless, tracking can be immediately restored by “predictive priming” of pyramidal cells at the place in tectum that represents the object after the saccade. This is achieved by TL transmitting two signals through SM to the pyramidal cell dendrites: (i) the pre-saccade position of the attentional focus and (ii) a corollary discharge giving saccade direction and amplitude. Only those pyramidal dendrites that bear the appropriate synapses decode the TL signals and prime the neuron by depolarizing it. The attentional locus now becomes established at the new tectal position and tracking resumes.

**Figure 7.**
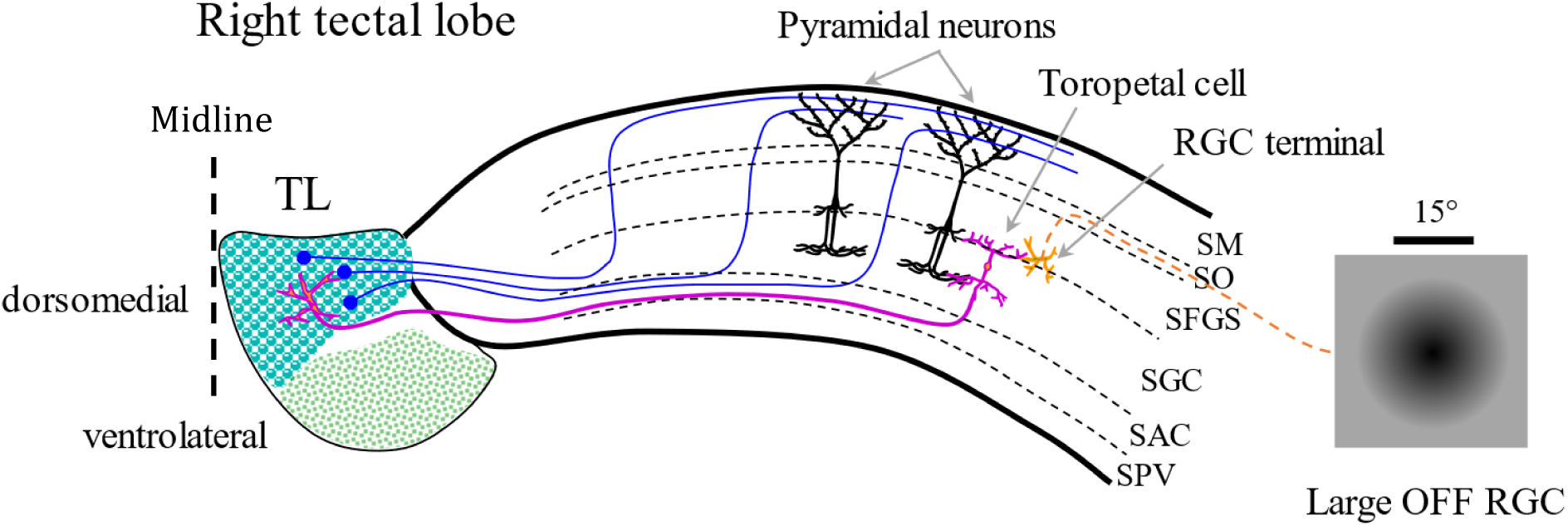
Coronal schematic of retino-tectal-TL circuitry. Retinal ganglion cells with large OFF receptive fields terminate in deep SFGS, synapsing excitatorily with Type X toropetal neurons which innervate dorsomedial TL. TL granule cells send marginal fibers through SM, synapsing excitatorily with the apical dendritic trees of Type I pyramidal cells. Depending on how toropetal inputs are switched in TL, different offsets in the priming of pyramidal cells can be achieved. This figure shows how RGC activation can lead to pyramidal cell priming at a higher position in the visual field.

**Figure 8.**
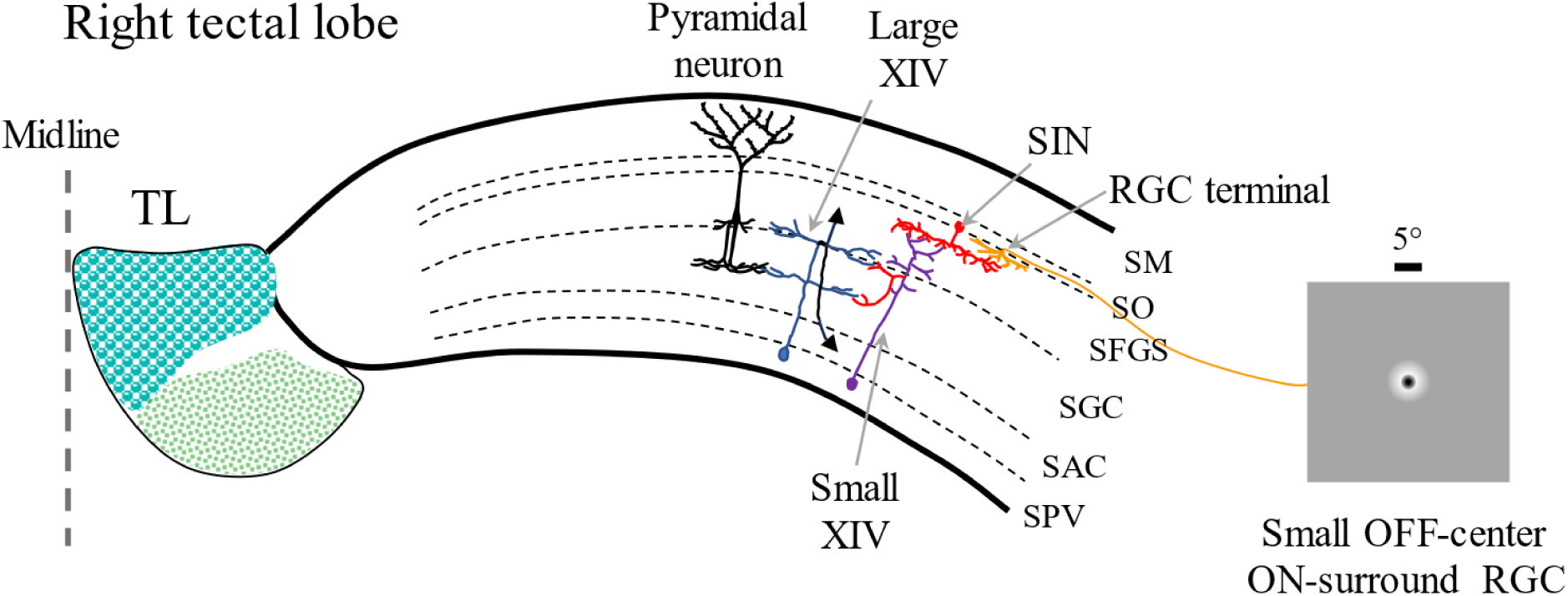
Coronal schematic of tectal circuitry. Retinal ganglion cells with small OFF-center, ON-surround receptive fields terminate in superficial SFGS, synapsing excitatorily with superficial inhibitory neurons (SIN) that inhibit Small Type XIV neurons; their axons inhibit neighboring Large Type XIV neurons which are excited by the axonal arbors of pyramidal cells. The output of the system is transmitted by the Large Type XIV axons that both ascend and descend (black arrows). Inhibitory processes are shown in red.

For the modelling of shape discrimination investigated here, the focus of attention, formed by heightened activity in a cluster of neighboring pyramidal neurons is supposed to select stimulus features that are close together in the retinal image, thereby “binding” them as a potential object. The focus of attention could then enable only the neural elements responding to those stimulus features, wherever the object is in the visual field. To obtain a reading of the object’s qualities, it is only necessary to collect or “bin” all the activity of the attentionally-enabled feature units across the tectal lobe. The resulting binned components provide a vector to identify the object, and to do it with positional invariance. The principle can be implemented with known tectal circuitry, while providing an explanation for how the basic visual feature information present in the superficial tectum could be used. However, this mechanism is unable to distinguish two objects with the same feature content but different spatial relationships between them, something that fish can clearly do (Sutherland & Bowman, 1969).

An aim of this paper is to show with the help of modelling, how neural structures in the midbrain of fishes could implement shape discrimination along these lines while improving on the simple binning mechanism. The most extensive studies of goldfish shape discrimination have been those of Stuart Sutherland and his associates undertaken in the development of his theory of pattern discrimination in animals and humans (Sutherland, 1968). Some of the data they generated by training goldfish on stimulus pairs and performing transfer tests with other pairs to uncover the features used for discrimination will provide tests of the model.

## Methods

In the Bowman & Sutherland (1969) experiments (“B&S” henceforward) the training and testing of goldfish used shapes cut from ¼” black plastic on a white background, each shape being separated horizontally by 5” (12.7 cm) and held at its center a small trough for a food pellet or a similar-looking inedible pebble. Each trial began with the fish swimming through an opening 12” (30.5 cms) from the shapes and swimming up to one of the shapes to receive either the food reward or the pebble, depending on the contingencies of the experiment. The decision point was not defined in these experiments but could have been between 30.5 and 6.5 cm, the latter distance set by the depth of a dividing screen projecting from the plane of the shapes.

The discriminations with my simulated neural system used 45 of the B&S stimulus shapes. They were scaled in size to present roughly the same area as the B&S stimuli (14.4 cm^2^). During a simulation run each shape was individually moved towards the camera from a starting distance of 19 cm up to 7.5 cm, representing changes in visual angle from about 11.4° to 28.1°, or a width increase by a factor of 2.46. The rate of approach was 0.3 pixels/tick over 200 simulation ticks. The starting positions of stimuli were randomized from trial to trial within a 32 x 24-pixel rectangle so that they activated different successions of pixels and receptive fields, leading to substantial trial-trial response variability.

### The Network

The methods for network construction and simulation are essentially those described for a model of tectum and TL (Northmore, 2017). A custom simulation program written in Borland Delphi Pascal allowed rectangular layers of artificial neural units to be arrayed in two dimensions of ‘‘brain space” and provided various methods for interconnecting them. Figure 2 shows the layers of the current network with the connections of a single representative unit in each of them. Figure 3 shows the activity within each layer at a moment in time during the approach of a 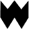 shape. Typically, each unit of a source layer projected synapses centered on the topographically corresponding point in the destination layer. Where the projection was spread over a region of the destination layer, the synaptic weights fell off from the center of projection according to a radial Gaussian function. A connection was specified by its maximum weight, either excitatory (+) or inhibitory (-), and by σ, its Gaussian spread in pixels (see values used in Fig. 2). Layers were composed of linear threshold units or of IAC units, the latter having been employed in Interactive and Competitive networks (McClelland & Rumelhart, 1988. See Northmore, 2017 and its supplemental data for details). An important feature of IAC units is that they obey 1^st^-order dynamics, in this case with a decay time constant of 10 simulation ticks. Consequently, an IAC unit in this network settles over time to a level of activation determined by the various excitatory and inhibitory inputs impinging upon it.

**Figure 2.**
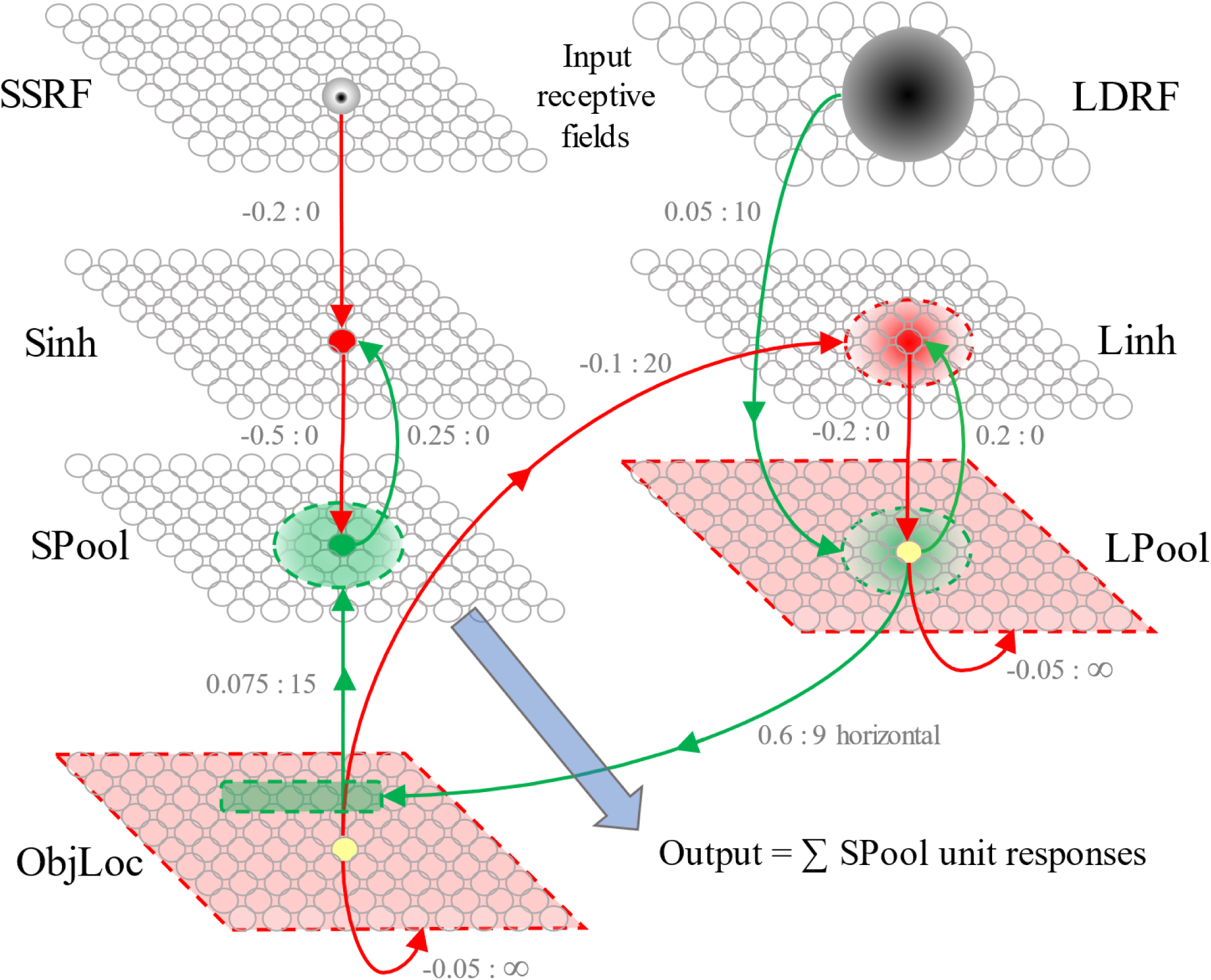
Schematic of the network. Each small colored disc represents an individual unit in its respective layer. Green arrows: excitatory synaptic connections; Red arrows: inhibitory synaptic connections. Arrows going to areas delimited by dashed lines show projective synaptic fields, either excitatory (light green), or inhibitory (light red), from the unit at the origin of the arrow. The two numbers next to each connection show its maximum weight and sigma of Gaussian spread.

**Figure 3.**
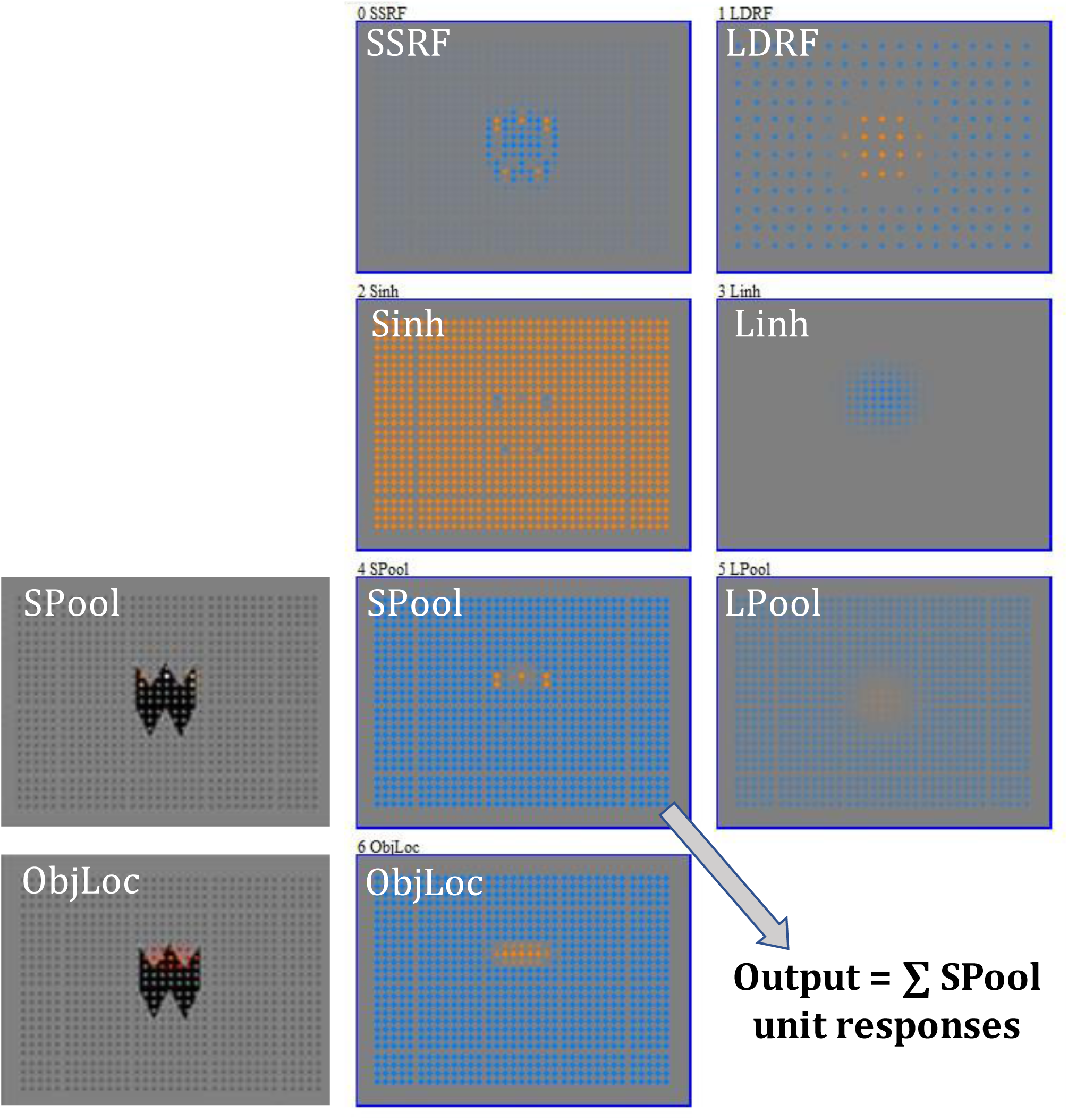
The network in action. The two rightmost columns show the layers currently processing the W shape during its approach. Orange is positive unit activation; blue is negative unit activation. Leftmost column: the W shape overlaid by unit activity; Upper: colored dots show SPool units responding to points on top; Lower: ObjLoc responses enabling SPool. Output of the model is the summation of all SPool unit responses.

The simulation program used GLscene, a 3-D graphics package based on OpenGL to present the black shape stimuli as sprites moving in 3D space against a gray background. The GLscene camera, representing the goldfish eye, viewed a rectangular 120 x 160-pixel array that was sampled by two kinds of difference-of-Gaussian receptive fields (RFs) emulating retinal ganglion cells (RGCs), and these constituted the visual input to the model. The smaller type, the small spot-detecting RF (SSRF) was designed to respond in a sustained fashion to the points and projections on the outlines of the black test shapes. It was composed of a central OFF Gaussian of 1.6° half-maximum diameter with a Gaussian ON surround of half-maximum diameter of 5.6° (see Figures 2, 7 & 8). The visual field was sampled by a 35 x 25 rectangular array of such units. The other field type, the large dimming RF (LDRF) was designed to give a sustained response to darkening within its receptive field with a half-maximum width of 15° but with no antagonistic surround (see Figures 2, 7 & 8). The visual field was sampled by a 17 x 12 rectangular array of such units.

In the previous simulation study (Northmore 2017), the layer of central importance was the “pointer network” whose activity formed an attentional focus for selecting incoming sensory information. It was identified with the population of pyramidal neurons in fish tectum. Here, the corresponding layer, composed of IAC units, is called the ObjLoc layer as it locates or points to the current object of interest in the visual field (see Figure 2). To implement the ObjLoc layer, each of its units should receive excitatory topographic visual input while making local connections that (i) inhibit all other units in the ObjLoc layer, thereby forming a winner-take-all circuit and (ii) excite units locally in the ObjLoc layer to allow the locus to move smoothly with movement of the attended object (for clarity not shown in Figure 2). The other layer type required for an attentional network are the “Pool” layers, also composed of IAC units and each subject to local inhibitory feedback from “inh” layers. In general, the Pool layers would receive and process a visual submodality (e.g. color, orientation, size) and interact with the ObjLoc layer, in a bidirectional fashion. In the current network, ObjLoc’s focus is activated from only one such pool, LPool, that indicates the center of dark shapes. ObjLoc’s focus enables the corresponding place in another pool, SPool, that responds to the pointed features of stimulus shapes. The model’s output is the activity of SPool under the focus of ObjLoc. Details of these layers and their connections follow.

In the network shown in Figure 2, the visual field is sampled by both the SSRFs and the LDRFs. Each LDRF connects excitatorily with Gaussian spread to LPool. Each LPool unit connects firstly, to excite a corresponding linear threshold unit in a layer of inhibitories, Linh, that connects back to it inhibitorily, and secondly, to inhibit all other LPool units.

Both of these connections limit the gain of this dimming pathway, focusing its activity pattern to roughly the center of gravity of a stimulus shape. Thirdly, each LPool unit projects excitatorily to the ObjLoc layer in a spatial pattern to bias the attentional focus in various ways. Three such patterns used are illustrated in Table 1 in green.

**Table 1.**
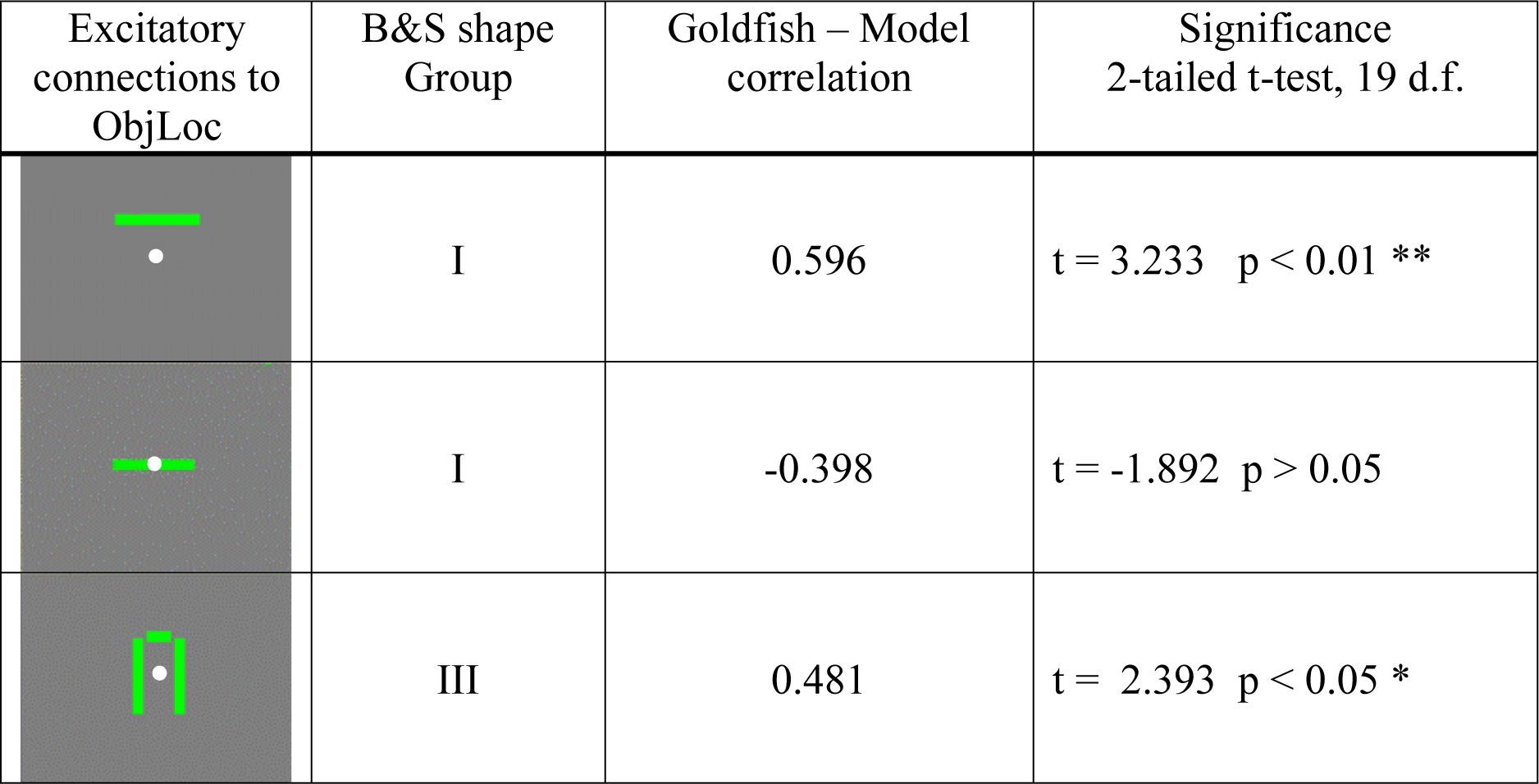
Three mappings of connections to ObjLoc from a single unit in LPool (left column, white disc). Each green bar represents a row or column of equally-weighted excitatory synapses onto the ObjLoc layer. The first row of the table shows a control condition without vertical offset that gave no significant correlation between goldfish transference and model response on Group I test pairs; the second with a vertical offset gave a significant positive correlation on Group I; the third with vertical and lateral offsets gave a significant correlation

| Excitatory connections to ObjLoc | B&S shape Group | Goldfish – Model correlation | Significance<br>2-tailed t-test, 19 d.f. |
| --- | --- | --- | --- |
| | I | 0.596 | $t = 3.233$ $p < 0.01$ ** |
| | I | -0.398 | $t = -1.892$ $p > 0.05$ |
| | III | 0.481 | $t = 2.393$ $p < 0.05$ * |

Upward biasing of attention was achieved by the 1^st^ row pattern in Table 1. A unit in LPool, exemplified by the white disc, makes excitatory connections (green) to a horizontal row (length = 9) of ObjLoc units, displaced to 3 rows above. The effect is shown in Fig. 1 where an M-shape is advanced toward the viewer while simultaneously shifting rightward. The LPool activity centroid tracks near the center of the stimulus shape while the ObjLoc activity centroid floats above. The LPool centroid rises a little above the center because it is in a positive feedback loop with ObjLoc via Linh. There is also some divergence of the two tracks as the image expands keeping the attentional focus hovering near the top.

**Figure 1.**
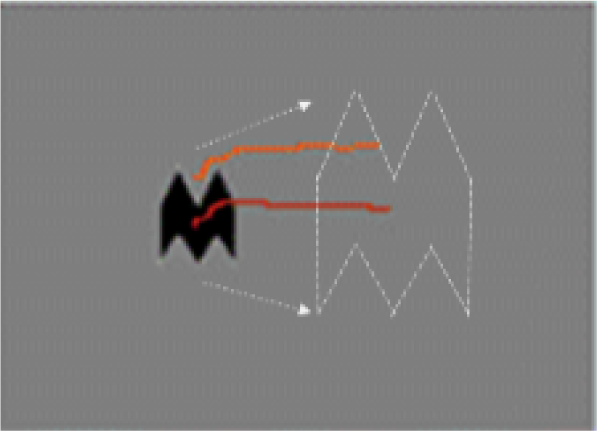
The M-shaped figure was moved rightward and toward the camera. The position of the centroid of LPool is shown in red; that of ObjLoc in orange.

The sharp points of shapes are responded to by the SSRFs. As Figure 2 shows, SSRFs inhibit their topographically corresponding linear threshold units in the layer Sinh, which in turn topographically inhibit units in SPool. Because the entire Sinh layer is biased on (see Fig. 3. Orange = +ve activation), SPool is generally inhibited (Blue = -ve activation), and any points of the stimulus shape leading to SSRF activation will disinhibit topographically corresponding units in SPool. However, the inhibition of SPool is too strong to allow the stimulus-dependent disinhibition alone to activate its units, but SPool units can be activated if they are also subject to excitatory drive from ObjLoc. Since this comes from a horizontal band of excitation (orange in ObjLoc in Figure 3) that hovers at the top of the shape, the SPool activation in total will depend on the number of points at the top. The final output of the network to the shape being analyzed is the summed response of all SPool units. It should be noted that if additional stimulus objects were to appear in the visual field they would be ignored by the network while ObjLoc is locked onto the attended stimulus.

Although the goldfish performing these visual discriminations would most probably use the front of its visual field, involving both eyes and both tectal lobes, I am making the simplifying assumption that each test shape is processed one at a time by one tectal lobe with a spatially uniform input from RGCs of the opposite eye. This is justified by the mapping data of Schwassmann & Kruger (1965) for goldfish.

## Results

To emulate the shape transfer experiments of B&S, each of their 45 individual black on gray shapes was input to the network by placing it randomly within a central rectangle (32 x 24 pixels) of the GLscene viewer and looming it toward the camera over the course of 200 simulation ticks, equivalent to a 2-sec approach time. Time was not a critical factor in the simulation as the decay time constants of the IAC units were short in comparison to the simulation period. The looming of each stimulus was repeated 20 times, generating waveforms for the summed responses of SPool such as those shown in Figure 4 **B**. The randomization of the stimulus starting positions and their subsequent trajectories introduced trial-to-trial variation in the network response as the receptive fields and network connections were activated differently on each trial. Varying stimulus position in this way was important for demonstrating position invariance of the model’s discrimination abilities claimed here.

**Figure 4.**
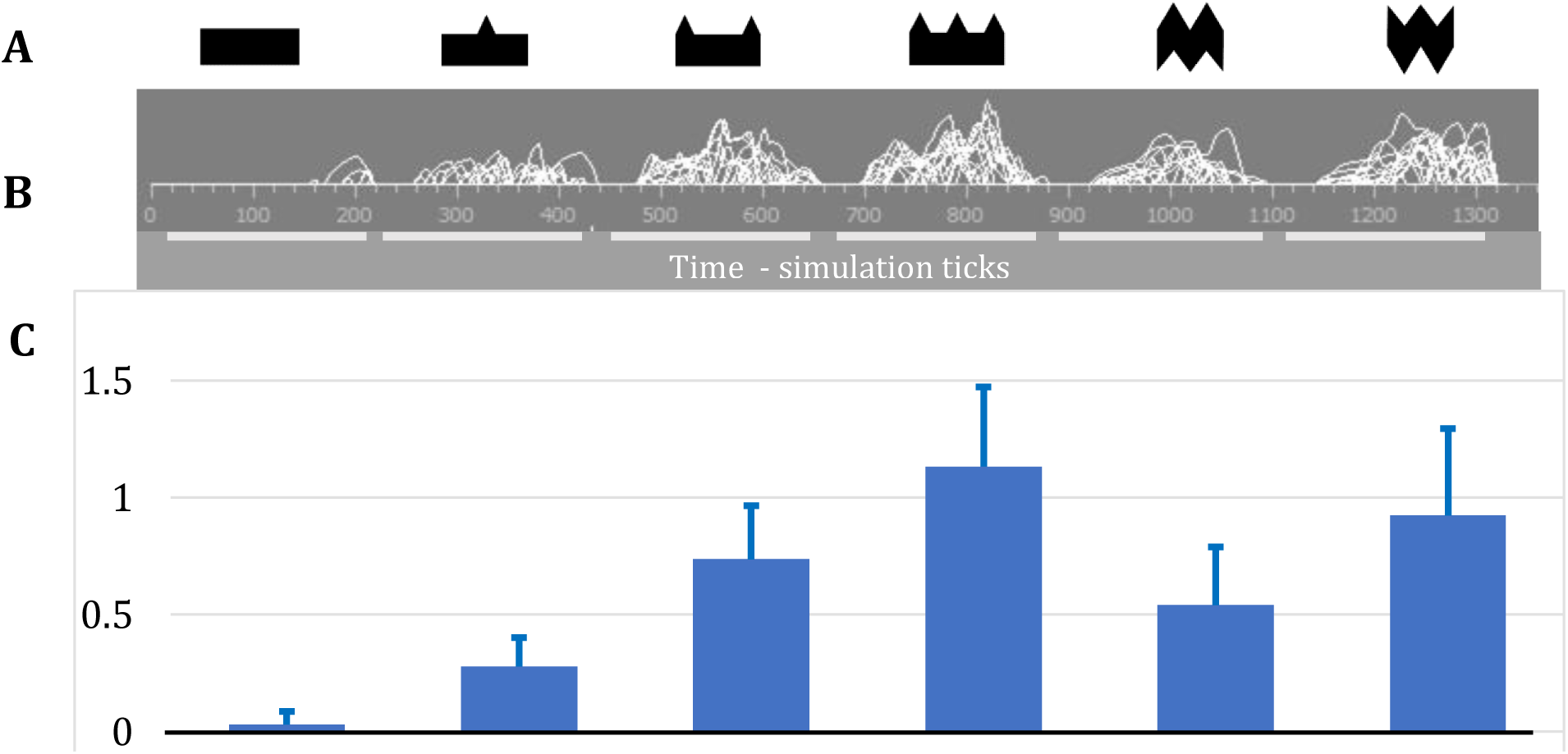
Network responseses derived from the summation of all SPool unit responses to six shapes (A) accumulated over 20 approaches (B), horizotal bars show movement duration. C Average of 20 responses integrated over time, with standard deviation bars.

Network responses to the individual shapes were obtained by averaging all the SPool unit responses over 20 approaches (see Figure 4 for examples). The model’s discrimination of a pair of shapes, one designated positive by B&S, the other negative, was assessed by subtracting the averaged SPool response to the negative shape from that to the positive shape, the difference giving the “Model Response” in figures 5 and 6. This value was compared with the corresponding goldfish “Transference” value for the stimulus pair using B&S’s percent correct (P), adjusted for 50% correct responding by chance i.e. (P-50)/50 (Abbott’s formula).

**Figure 5.**
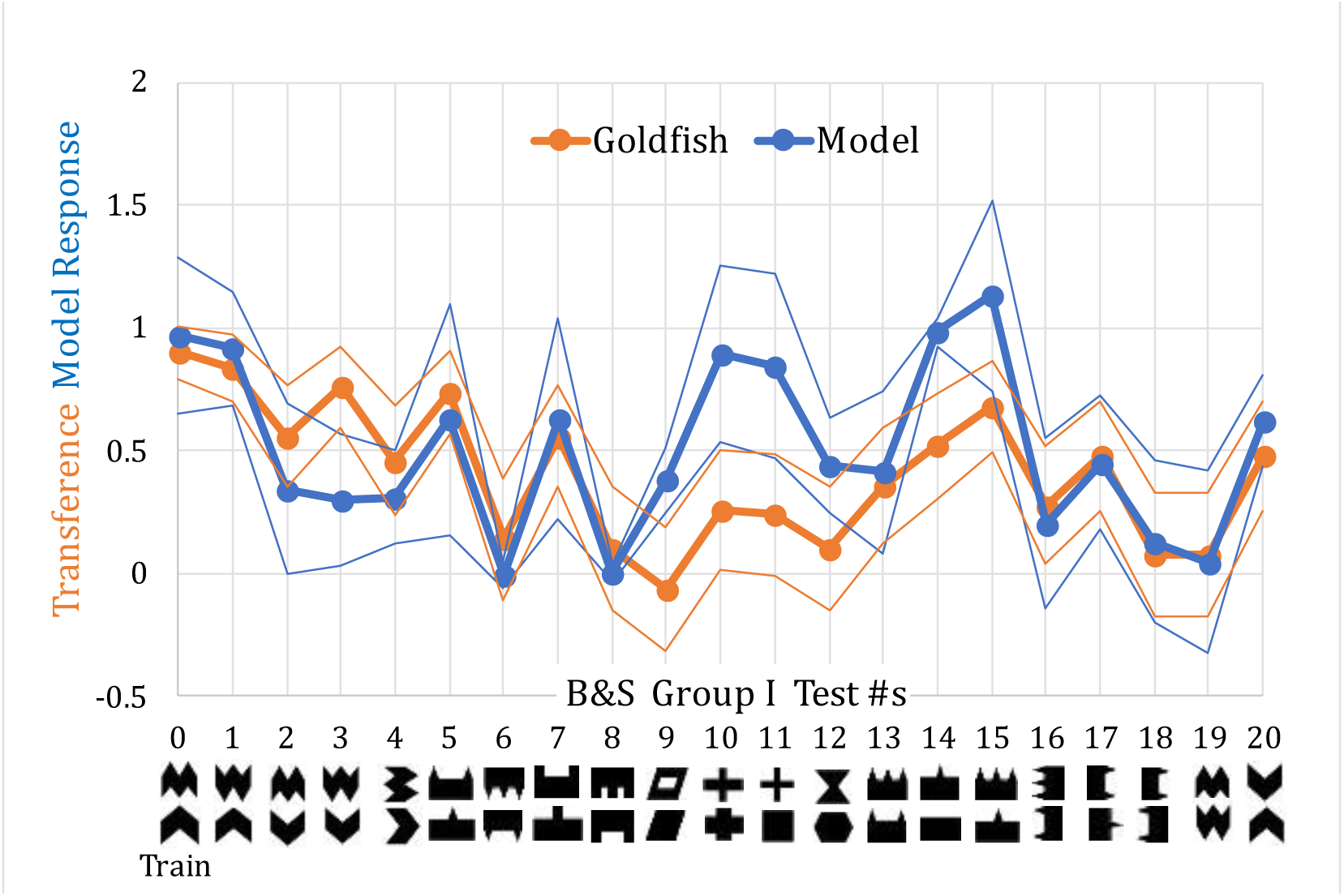
Bowman & Sutherland’s Group I goldfish behavioral transference (orange) compared to the model response (blue). Fish were trained to 95% correct on the leftmost test pair and tested on the other 19 pairs. The goldfish data and 95% confidence intervals from the binomial distribution based on 60 trials were corrected for 50% chance responding by Abbott’s formula. The model data points and 95% confidence intervals were based on 20 trials each.

**Figure 6.**
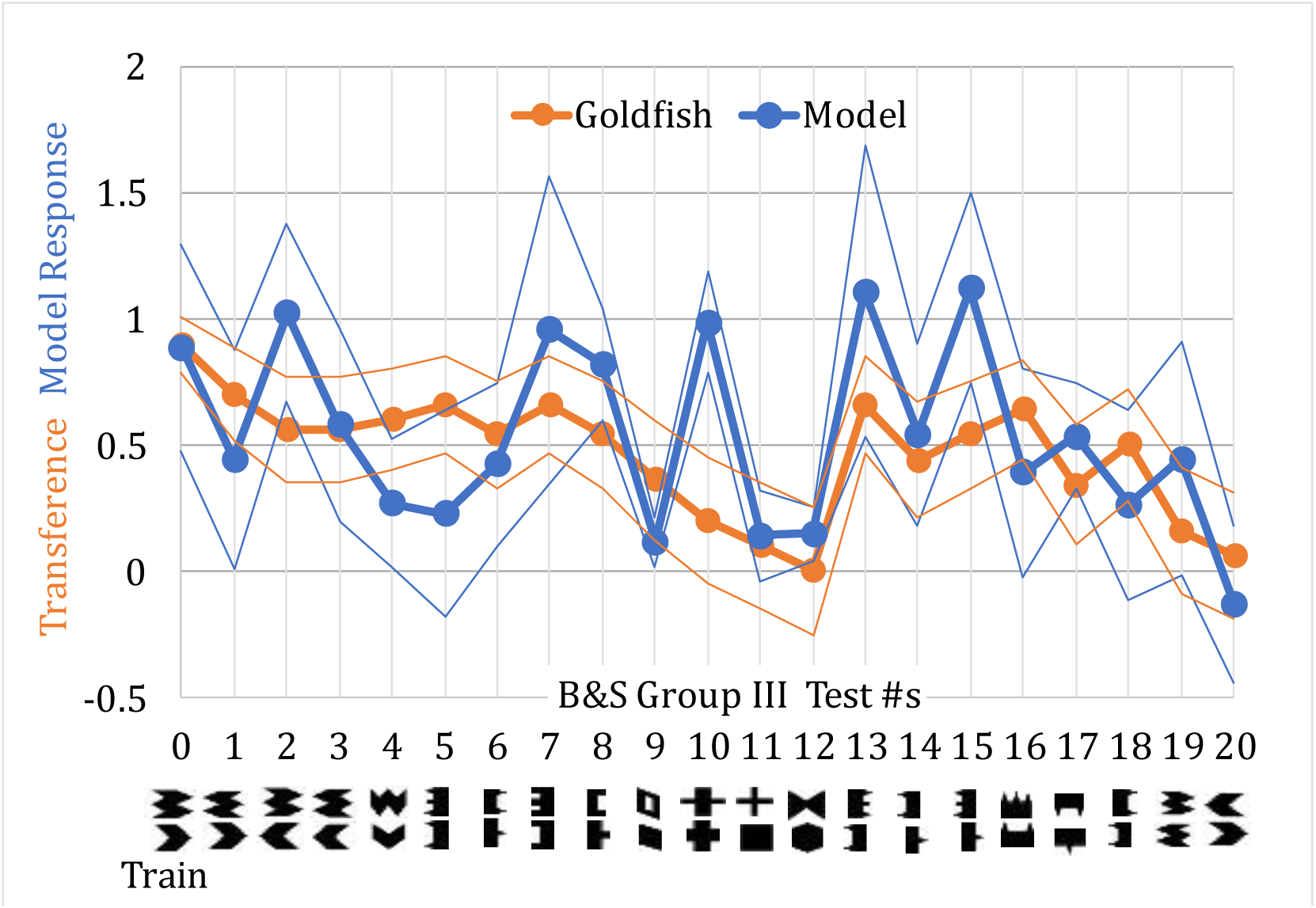
B &S’s Group III goldfish behavioral transference (orange) compared to the model response (blue). Details as for Fig. 5.

Transference, so defined, varied from about 0 to 1.0 making it comparable to the range of Model Responses, both values being plotted in figures 5 and 6.

### Upward biasing for Group I

In B&S’s Group I test, fish were trained to discriminate 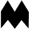 from 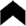 (leftmost pair in Fig. 5), giving the fish the opportunity to use 2 vs.1 points on top, and/or 3 vs 2 points on the bottom. Figure 5 presents the goldfish data (orange) for the other 20 pairs showing that transference was highest when there was a preponderance of points on top of the positive shape (upper in each pair). B&S concluded that transfer was governed by the relative numbers of points on top, not the absolute number.

This tendency of goldfish to attend to the uppermost discontinuities of shape outlines requires a means of locating the tops. The method employed by the network of Fig 2 is to locate the center of darkness in the shape using an array of large OFF receptive fields, and send its position to ObjLoc with an upward offset (Table 1, 1st row).

Figure 5 presents the model response (blue) for all 21 pairs in the B&S Group I test. Many of the high and low points of the goldfish performance are paralleled by the model response. Thus, ‘positive’ shapes with more points on top than their negative counterparts yielded high responses, whereas those with few or no points on top yielded the lowest responses. The greatest discrepancies, where the model responded excessively, were for positive shapes with pronounced vertical projections like the crosses (tests 10 & 11) and shapes with a preponderance of upward projections (tests 14 & 15). A significant correlation (r=0.596) between goldfish transference and model response was obtained (Table 1, 1st row). To show that the vertical offset was critical to the model’s emulation, a control condition with no offset was run, yielding no significant correlation (Table 1, 2nd row).

### Left, right and upward biasing for Group III

In the Group I experiment of B&S, transference to shapes with points on the sides was poor, in keeping with the goldfish’s natural tendency to focus attention on the tops of figures. In their Group III experiment, fish were trained on the pair of shapes 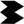 and 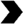 with points on both left and right sides in the ratios 3:2 and 2:1 (Figure 6 leftmost pair). Fish successfully achieved criterion discrimination (95%) and were tested on all the other pairs shown in Figure 6. They showed good transfer to the “pointier” shape pairs (tests 1-3 & 5-8) whether the points were on the left or right, but little or no transfer to “pointless” shapes (tests 9-12), nor to pairs of mirror-image “pointed” shapes (tests 19 & 20). They transferred well in test 4, a pair that differed in the number of points on top, presumably reflecting the natural preference.

To show that the model can also emulate these data, the connections from each LPool unit to ObjLoc were made to be columns of excitatory synapses 10 units high biased 3 units leftward and rightward, and a row of synapses 3 units long and biased upward by 5 units to form the pattern shown in Table 1, 3^rd^ row. Each individual shape of Figure 6 was loomed to the camera to collect the integrated responses of SPool units as before. The integrated responses to the positive and negative shapes of each test pair were differenced to give the Model Response, which was plotted with the goldfish Transference in Figure 6.

The model generally reproduced the overall profile of the goldfish transference, albeit in more exaggerated form, yielding a significant correlation of 0.481 (Table 1, 3^rd^ row). The only significant discrepancy was the high response of the model to the two-crosses pair (test 10); the thin cross bars projecting from a thick body strongly activated the point detectors, more so than they did when projecting from a thin body (test 11). The reason is that the overall black area of the latter cross was relatively small, leading to a weaker drive of LPool to ObjLoc and weaker point detection. Eliminating test 10 gave a higher correlation (r = 0.593, p < 0.01).

## Discussion

B&S concluded that the main determinant of goldfish discrimination performance was the relative number of points at the tops of shapes (see also Sutherland & Bowman, 1969), rather than the absolute numbers of points on the training shapes. The model similarly responds to relative numbers because of its simple decision rule: treat the shape with the higher SPool activation as the positive stimulus. The model, following this rule, also accounted for goldfish discrimination performance when trained to attend to left, right and top edges of shapes.

### Neural implementation

Here I attempt to link the structure and connectivity of the network with the neurology of the visual system in fishes.

The small and large receptive field types, SSRF and LDRF, providing visual input to the model, have their counterparts among goldfish retinal ganglion cells. LDRF is a sustained OFF receptive field without antagonistic surround subtending 15°. These resemble alpha-like retinal ganglion cells in goldfish (Cook et al., 1992), that exhibit sustained OFF-firing and subtend 13°-24° under light adapted conditions (Northmore & Oh, 1998). SSRF, the OFF-center ON-surround receptive field had an overall half-width of 5.6°, close to the smallest RGCs of goldfish (Northmore, 1977). Its spatial frequency response peaked at 0.33 cycles/degree, which is also where the behavioral contrast sensitivity function of goldfish peaks (Northmore & Dvorak, 1979), corresponding to a point-spread function with a center subtending 1.6-1.7°, which matches the OFF-center half-width of SSRF.

Both electrical recordings and Ca^++^ imaging from superficial tectum show further processing of retinal inputs, elaborating specificity for size, orientation, direction of motion and color (Gabriel et al., 2012; Maxsimova et al., 2019; Nikolaou et al., 2012; Nilo et al., 2021). These provide the ingredients for the binning method of object recognition, whereby the object being attended is specified by the amounts of the different features present within the current focus of attention. However, the present simulation results suggest that recognition of black shapes can be largely accounted for with only one type of feature detector, the SSRFs, that are selective for small dark points.

LDRFs, in a relatively coarse array, connect excitatorily and topographically to LPool, a layer of units that can be identified with the Type X neurons of Meek & Schellart (1978), also known as toropetal neurons because they project to dorsomedial TL (Xue et al., 2003; Folgueira et al., 2007) and are responsible for TL’s retinotopic dimming activity (Northmore, 1984; Robles et al., 2021). The dendrites of toropetal neurons in goldfish ramify in deep SFGS and mid *stratum griseum centrale* (SGC), which is where the axons of dimming RGCs terminate (Robles et al., 2021) (see Figure 7). It is also noteworthy that the toropetals ramify at the same depth as (i) the pyramidal neurons’ soma and near-soma dendrites and (ii) the deep axonal and dendritic ramifications of pyramidal neurons (Xue et al., 2003; Folgueira et al., 2020). The close relationship between these two neuron types suggests exchanges of information can take place between them such as were previously postulated by Northmore (2017) as part of the mechanism to maintain attention across saccadic eye movements. In the present network, the connections of special interest, those from the toropetals (LPool) to the pyramidal neurons (ObjLoc) are the excitatory, displaced connections (Table 1; Figure 2). The key idea here is that the displacement is achieved at the final arrow in the pathway: toropetal → dorsomedial TL → marginal fibers → pyramidal neuron dendrites (see Figure 7). The position of the center of the stimulus shape is represented by an activity peak on the layer of toropetals (LPool) and is sharpened by local and lateral feedback inhibitory connections between Linh and LPool. The toropetals, where their activity peaks, excite neurons in dorsomedial TL at the topographically corresponding position along TL’s length. There, TL neurons transmit excitation through SM in a set of fibers that preferentially excite the dendrites of pyramidal neurons at a more dorso-medial position in tectum than the position of peak activity in the toropetal layer, i.e. corresponding to a greater elevation in the visual field. This is illustrated in Fig, 7 by a second, more medially situated pyramidal neuron.

According to the model, the neural circuitry determining the center of a shape is driven by a dimming pathway. It is interesting to note that Mackintosh & Sutherland (1963) reported that black on white shapes were much easier for goldfish to learn than white on black, evidence of a special role of dimming pathways whose characteristics and origins have been discussed elsewhere (Northmore, 2023a).

A neural scheme for the point-detecting circuitry of the model (vertical pathways on left side of Fig. 2) is suggested in Figure 8. Small OFF-center, ON-surround RGCs make terminal arbors in a superficial sublayer of SFGS. The sign reversal of the retinal input onto the inhibitory Sinh layer could be achieved if the retinal terminals were to excite the superficial inhibitory interneurons (SINs) that ramify in these superficial layers (Preuss et al., 2014), which in turn inhibit the neurons corresponding to the Sinh layer. A putative assignment of Sinh units would be to the small Type XIV neurons of Meek & Schellart (1978) which have short processes at different depths in SFGS and SGC. If these are equivalent to the non-stratified periventricular neurons of zebrafish larvae (Robles et al., 2011), their branches in SFGS would be dendrites, contacted and inhibited by the SINs; while their axons would be GABAergic, inhibiting the neurons composing the SPool layer. Small Type XIV neurons in goldfish were identified and recorded by Guthrie & Sharma (1991) who found them to be tonically active, but modulated by visual stimulation. They could, therefore provide continuous inhibition of Spool units while conveying stimulus feature information to Spool as required by the model. These small Type XIV likely arise from the numerous GABAergic perikarya in SPV (Médina et al., 1994; Maruska et al., 2016). SPool neurons are proposed to be large Type XIV neurons of Meek & Schellart (1978) whose dendrites ramify at the SFGS/SGC border, and in the middle of SGC. Either or both of these dendrites could be the targets of inhibition from the small Type XIV. The deeper branches of the large Type XIV, those in mid SGC, are positioned to receive input from the pyramidal neurons (ObjLoc) which the model assumes to be excitatory, and most probably, glutamatergic (Kageyama & Meyer, 1989). These large Type XIV also have axons, usually branching into two, one of which exits tectum (Kinoshita et al., 2006). It is these axons that in the aggregate, form the output of the model, supplying information about the preponderance of different features in the attended object.

Next, the system needs to collect this feature preponderance in both lobes of tectum for deciding which shape to approach. There should also be separate short-term memory storage of feature preponderance pertaining to the stimulus shapes over the few seconds it takes the fish to make its choice, accumulating information about the two shapes until one of them wins out. These processes, while possibly involving extra-tectal centers, are likely anchored to the tectal topographic map, initiating and directing approach to the chosen shape via the set of labelled lines from different tectal regions to premotor centers in the brainstem commanding turning movements of specific amplitudes (Helmbrecht et al., 2018). The model suggests that it is the locus of attention on the tectal map that directs approach and this is borne out by the observation of B&S that goldfish snapped at the tops or at the sides of the stimulus shapes, depending on their training.

### What is being learned?

Sutherland’s (1968) theory held that learning a pattern discrimination first involves learning to “switch in” stimulus analyzers that are suitably discriminative for the task, and secondly to learn to associate those analyzers with an appropriate behavioral response, such as approach or avoid. The present model has some bearing on the first stage. I chose what seemed to be a successful type of analyzer for discriminating the B&S shapes, namely the SSRFs, but in reality, a number of basic analyzers for size and orientation are available from neuronal elements in superficial tectum. To use these for discrimination is simple in principle. Instead of accumulating activity in SPool only, accumulations in separate pools for different line orientations, stimulus sizes and colors would effectively generate a feature vector for the attentionally selected stimulus object. If approach to this object leads to reward, a Hebbian-type pattern associator could adjust the weight of each vector component to raise approach probability on a later trial. Thus, appropriate analyzers are “switched in.”

The mechanism just sketched, in effect, throws all of an attended object’s features into one bin; features of a different object, attended to later, will go into another bin, allowing the two to be discriminated. This process would have been available to primitive creatures given the kind of winner-take-all visual attention modelled here. An obvious limitation to the binning principle is that the feature accumulation takes place without regard to the spatial relationships between features.

### TL as an attentional lens

The ray-fined fishes, the actinopterygians, when they emerged some 400 million years ago developed a tectal add-on, the TL (Yamamoto & Hagio, 2021), which I maintain allows visual attention to be deflected in a controlled fashion. The predictive priming model for holding attention on a prey object in the face of eye saccades is an example (Northmore, 2017). That model envisioned TL converging two kinds of information onto the tectal pyramidal neurons: the pre-saccade position of the object, and the upcoming saccade vector. The two together could be decoded by a learned pattern of synapses on pyramidal neuron dendritic trees leading to the priming of only those pyramidal neurons at the predicted post-saccade position. Combining the spread of pyramidal dendrites and the spread of a TL neuron’s axons across tectum, which may be considerable (DeMarco et al., 2021), the system could perform shifts of up to 30° of visual angle, the maximum amplitude of goldfish saccades.

The current model also envisions displacements of the focus of attention, in this case for selecting features from parts of an object. Binning these allows discriminations between many more patterns. The B&S experiments and the emulation of them have shown that fish can displace attention not only upwards, as they seem to do naturally, but also to both sides, and to do so simultaneously, representing a diffusion of attention. If the predictive priming theory is any guide, displacements and diffusions of up to 30° may be possible, allowing a good measure of constancy in the assessment of features of retinal images of different sizes. Thus, TL seems to be an adjustable refractor of the spotlight of attention.

To achieve control over TL’s refractions, one might assume an arrangement analogous to saccadic predictive priming (Northmore, 2017) in which TL transmits two kinds of signals in SM to achieve attentional shifts. As in the prior model, the first type of signal represents the position of the object’s center derived from its topographically mapped tectal activity relayed via toropetals to TL. There, granule cells distribute activity through marginal fibers in SM over a wide swathe of tectum centered on the object’s representation. Instead of the saccadic corollary discharge of the prior model, the second type of signal transmitted by TL, is a discrimination-task-related control signal generated along the length of TL specifying the direction and extent to which the tectal locus of attention should be deflected. These two TL signals are decoded by patterns of synapses between marginal fibers and the dendritic trees of the pyramidal cells such that the attentional locus of pyramidal cell layer activity is shifted or spread. The source of the task-related control signals in TL can be surmised to descend from the telencephalon via pretectal nuclei (Wulliman, 1994; Yañez et al., 2018; Xue et al., 2003; Folgueira et al., 2020; Yamamoto & Hagio, 2021). Learning a discrimination, therefore, not only requires switching in the right feature analyzers, but also controlling TL to make attentional refractions for the task at hand.

### Experimental support for the model

One of the key assumptions made by the model is that pyramidal neurons of tectum form an attentional focus of heightened activity that selectively highlights a visual stimulus. It may be an individual pyramidal neuron or a local group that is involved. The focus is assumed to be dynamic in being able to jump from one stimulus to another and to be able to track a moving stimulus in space. The focus, when activated, selects stimulus features for analysis, memorization, and control of behavior. As yet, there is no direct evidence for pyramidal cells forming an attentional locus but selective imaging of pyramidal cell activity in larval zebrafish has already been achieved (Tesmer et al., 2022) and could be used to look for it. So far, the best supporting evidence is the demonstration by Romano et al. (2015) in larval zebrafish tectum that the neuronal response to two simultaneous light spots was less than that to the spots individually, demonstrating a widespread mutual inhibitory interaction across tectum.

Another proposal of the model is that TL and the marginal fibers provide a mechanism for fine-tuning or refracting the spotlight of attention, to improve pattern discrimination. Ablation of TL in goldfish produces no obvious behavioral effects except for a reduction of the dorsal light reflex (Gibbs & Northmore, 1996), perhaps by disrupting dimming pathways. According to the model, however, TL ablation would be expected to affect pattern discriminations and the learning of them. A series of experiments by Mark and associates (Mark, 1966; Mark et al., 1973) on a cichlid showed that sectioning the tectal commissure overlying TL prevents the interocular transfer of learned brightness, color, and pattern discriminations. Although the commissure and the TL are contiguous, their relationship is unclear, but it is likely, as Mark admits, that damage was done to TL, especially while separating the two halves of TL at its anterior end. Tesmer et al. (2022) showed in larval zebrafish that TL on one side receives visual input from toropetal neurons in its contralateral as well as its ipsilateral tectal lobe, allowing pyramidal cells to be influence by either eye (see also Northmore 2023b). Mark’s midline sectioning with the resulting failure of interocular transfer of learned discriminations may be explained, at least in part, by the interruption of TL control signals. That sectioning affected discriminations of brightness and color suggests that TL might exert control over more than just shape features.

In another cichlid experiment, Mark et al. (1973) scraped the surface of tectum to ablate the marginal layer. This produced no gross blindness in fish but a transient impairment in discrimination learning. Interestingly, they observed a very similar transient impairment after complete forebrain ablation, consistent with the suggestion already made that TL control signals descend from forebrain via pretectum.

### Implications for theories of pattern recognition

It is interesting to consider how the present modelling effort bears upon Sutherland’s (1968) theory of pattern recognition, in particular, whether it has solved any of the 12 conditions that he stipulated for a satisfactory theory. Half of his conditions have been satisfied successfully by the model, the remainder being beyond its scope. His *Position Invariance* is well handled by the binning of features at the attentional locus, as is *Size Invariance*, at least up to an extent determined by TL’s spread of priming capacity. *Brightness Invariance*, or more precisely contrast-reversal invariance, could be handled with receptive fields with reverse contrast preferences. *Lack of Invariance under most Rotations* and *Confusions Made between Patterns*, in certain instances, could be explained by the settings of TL-mediated biases, for example to tops but not to sides of patterns. Lastly, his *Consistency with Physiological Knowledge* has been the watchword of this study.

The thrust of Sutherland’s theory was that the output of the feature processors leads to the writing of descriptive rules for shapes into a store during learning, and that the outcomes of subsequent testing depend on how these outputs match the descriptive rules. What neurally inspired modelling has shown here, as in other instances, is how rules can, and perhaps should be superseded by networks (Rumelhart & McClelland, 1986).

## Abbreviations

B&S: Bowman & Sutherland (1969)
SM: *Stratum marginale*
SO: *stratum opticum*
SFGS: *stratum fibrosum et griseum superficiale*
SGC: *stratum griseum centrale*
SAC: *stratum album centrale*
SPV: *stratum periventriculare*
TL: *torus longitudinalis*
RF: receptive field
RGC: retinal ganglion cell
SIN: superficial inhibitory neuron
SSRF: small spot-detecting receptive field
LDRF: large dimming receptive field
IAC: interactive and competitive
GABA: gamma amino butyric acid
mya: million years ago.

